# Linking plantain derived metabolites in sheep urine with nitrification inhibition in soil

**DOI:** 10.64898/2026.07.01.735958

**Authors:** Michelle Peterson, Nigel Joyce, John van Klink, Glenn Judson, Trish Fraser, Craig Anderson

## Abstract

Metabolites from *Plantago lanceolata* (plantain) biomass have been linked with biological nitrification inhibition (BNI) in soil. After grazing, leaf metabolite chemistry is altered via digestion, and a suite of secondary metabolites are then delivered onto soil via dung and urine. The purpose of this study was to establish if urine from sheep grazed on plantain had BNI activity when added to pasture soil, and to identify the metabolite profile(s) that most likely contribute to the BNI effects observed. Groups of sheep (n=5 sheep per group) were grazed on one of nine different plantain cultivars in autumn and spring with analysis of leaf material, urine, soil incubation and BNI bioassay data used to identify potential metabolite candidates implicated with BNI. The urinary nitrogen and metabolite composition of sheep fed plantain varied significantly between cultivars and season. After 28 days of incubation, all soil microcosms treated with plantain-derived urine had up to 35% less nitrate than comparative ryegrass urine controls in both seasons, except one in autumn. The key phytochemistry associated with lower soil nitrate concentrations was phenylethanoid and iridoid glycosides resulting in a higher output of glucuronidated, methylated and sulfated secondary metabolites in the urine. Among 19 secondary metabolites identified in the urine, hydroxytyrosol-related metabolites as well as catechol glucuronide, 2-methoxyphenyl sulfate and guaiacol-ß-D-glucuronide appear to be the most likely target compounds with respect to the BNI effects observed. Variation in metabolites from different plantain cultivars affected the ratio of metabolite derivatives in urine, which ultimately affected soil nitrification rates. Cultivar phytochemistry is therefore an important consideration with respect to BNI under urine patches.

**Highlights:**

- Sheep grazing different plantain cultivars had different urine compositions
- Urines elicited biological nitrification inhibition (BNI) in soil and *in vitro*
- Different BNI response was related to differential expression of urine metabolites
- Key urine metabolites associated with BNI are derived from glycosidic compounds

---

Keywords: *Plantago lanceolata*, plantain, soil, ovine urine, biological nitrification inhibition, plant secondary metabolites.

## 1. Introduction

The genus *Plantago* (Plantaginaceae) is known to contain a suite of plant secondary metabolites (PSM) including the iridoid glycosides (IrG) aucubin and catalpol, and the phenylpropanoid glycoside (PhG) verbascoside (syn: acteoside). Within a human health context, these compounds have been identified as having a wide range of pharmacological activities, including antimicrobial, anti-inflammatory, antioxidative, hepatoprotective and antihyperglycaemic properties (Bhattamisra et al., 2019; Zeng et al., 2020; Xiao et al., 2022).

Ribwort plantain (*Plantago lanceolata* L.), a herb commonly found in New Zealand pastures, has relatively high concentrations of aucubin and verbascoside (up to 3 and 9%, respectively), although this is dependent on cultivar, season, temperature, shading, plant age and nutrient supply (Adler et al., 1995; Tamura, 2001; Tamura, 2002; Tamura and Nishibe, 2002; Al-Mamun et al., 2008; Navarrete et al., 2016; Box and Judson, 2018; Box et al., 2019). Studies in the field, and in laboratory incubations, have found evidence for reduced nitrogen (N) mineralisation and nitrification in soil when ribwort plantain was included in the sward or when under-seeded with potatoes (Rauber et al., 2008; Dietz et al., 2013; Massaccesi et al., 2015; Carlton et al., 2019; Peterson et al., 2023). In the absence of any other explanation, it has been suggested that plantain PSM (or metabolites hereafter) exuded from the roots act as inhibitors of soil nitrification by interfering with the biological oxidation of ammonium to leachable nitrate via the intermediary nitrite. Inclusion of ribwort plantain in pastoral swards is therefore considered to be a potential avenue for reducing nitrate leaching from animal farming systems. Changes in nitrate concentrations and N-cycling dynamics may also provide an avenue for reducing nitrous oxide emissions as less nitrate is available to be denitrified (Rodriguez et al., 2023). This would be valuable for all food production systems for optimal N-use efficiency.

Alongside ‘direct’ biological nitrification inhibition (BNI) via root exudation and diluted urinary-N through the diuretic effect of plantain consumption, there is evidence for an ‘indirect’ BNI effect by compounds excreted in the urine of ruminants that consume plantain. Urine metabolite profiles are known to depend on plant species and the complement of plant metabolites ingested (Keir et al., 2001; Lane et al., 2006). Metabolites excreted in ruminant urine that are thought to be derived from the consumption of plantain (cultivar ‘Agritonic’) have been implicated in the reduction of net nitrification in laboratory soil microcosms compared with ryegrass (*Lolium perenne* L) (Judson et al., 2018; Judson et al., 2019). In microcosms treated with urine from sheep or from cattle fed plantain, a delay in nitrate production in soil of up to 14 days was observed when compared with no-urine controls. Beyond day 14, nitrate production accelerated but at no point reached the concentrations observed in microcosms treated with urine from ruminants consuming ryegrass.

A definitive correlation between nitrification inhibition in soil, and the relative abundance of specific metabolites that might be implicated in BNI (such as aucubin, catalpol and verbascoside) has not yet been made. However, there is evidence that these metabolites are involved, including lowering N excretion from animals (Navarrete et al., 2016; Gardiner et al., 2020; Peterson et al., 2023). Peterson et al. (2022) reported 800 mass features detected by liquid chromatography combined with mass spectrometry (LC-MS) in the urine of sheep fed plantain that were unique or present in much greater concentrations than in the urine from sheep that were fed ryegrass. Nitrogen-containing metabolites, defined as ‘non-urea nitrogenous compounds’ (NUNCs) within the urinary organic fraction (UOF) were twice as numerous in plantain urine than ryegrass urine and many looked to be sulfated, methylated and glycine conjugates (Peterson et al., 2022). Verbascoside, aucubin and catalpol were not identified in the urine, but possible derivatives of these metabolites were. Subsequently, a tentative linkage between nitrification inhibition and phenolic compounds derived from urine was made (Peterson et al., 2022).

This study aimed to confirm the identity and postulate roles of key plantain-derived urine metabolites with respect to BNI in soil. It was hypothesised that variable expression of shared metabolite commonalities between plantain cultivars would result in different ratios of inhibitory metabolites carried through to the urine, thus conferring different impacts on nitrification inhibition in the soil. The inhibitory capacity of the urine was assessed both in soil and *in vitro* bioassays, with LC-MS conducted on both plant and urine samples to find metabolite commonalities shared across plantain cultivars that correlated with the reductions in nitrification rates observed.

## 2. Materials and methods

A flow diagram summarising the experimental design, analysis types, replicate and sub-sample structure is presented in Figure 1.

**Figure 1:**
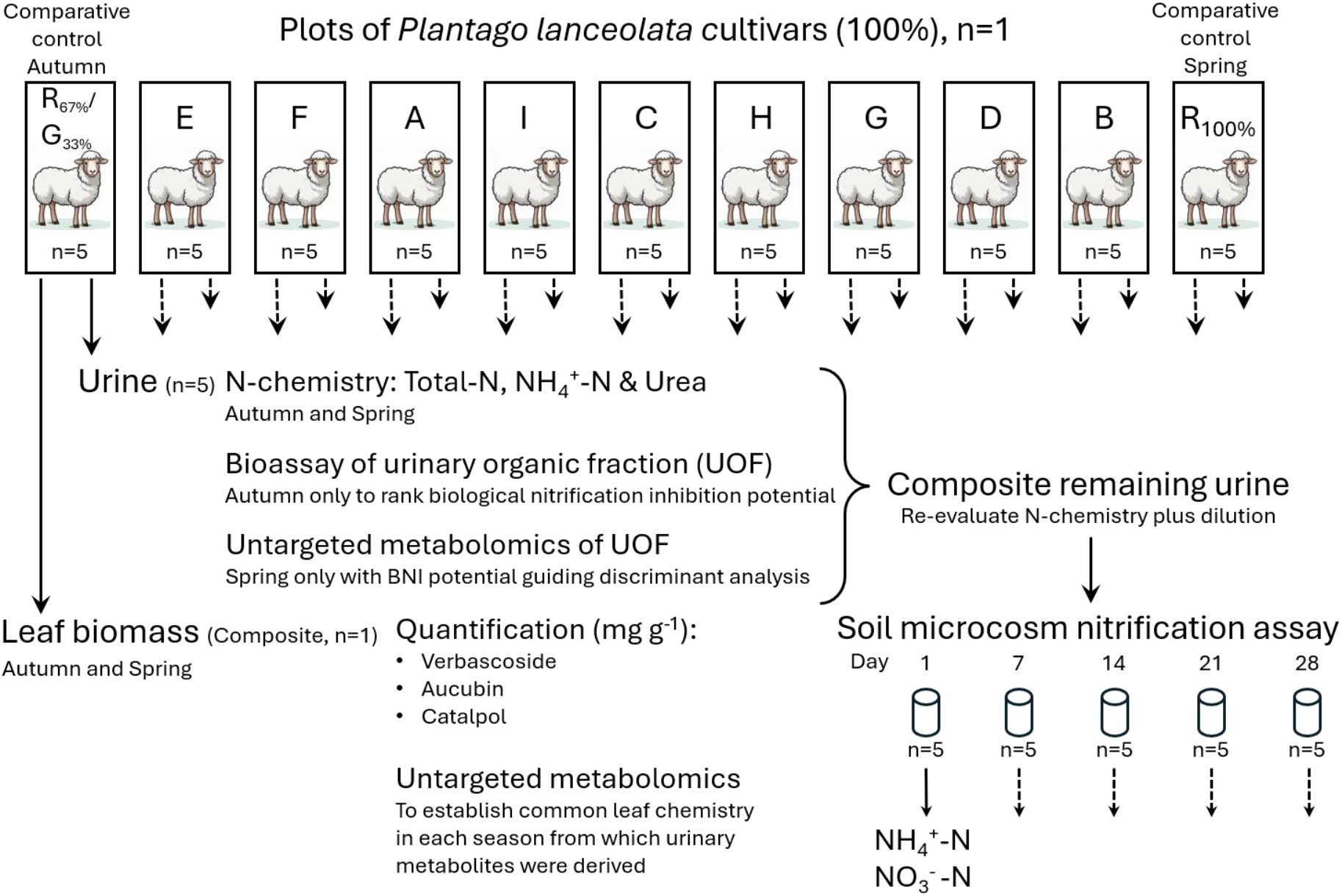
Flow diagram illustrating trial and experimental structure. The trial had singular large plots of nine different *Plantago lanceolata* cultivars designated A to I. Two different controls were used depending on season, with urine sourced from animals in an indoor feeding trial used in autumn (fed a mixture of ryegrass, 67%, and Cultivar G, 33%), while in spring urine came from animals grazing an adjacent 100% ryegrass plot. Each plot was grazed at a stocking rate of 5 sheep per 2500 m^2^ plot area. When sampling occurred, urine from each animal was collected separately, and a composite leaf biomass sample was taken from each plot. Urinary N chemistry and the nitrification inhibition capacity of the urinary organic fraction (via bioassay) were measured on subsamples for each individual sheep in autumn. In spring, urinary N chemistry along with the metabolome of the urinary organic fraction was measured on subsamples for each individual sheep. After subsampling for chemical, bioassay and metabolome analysis, remaining urine from sheep grazed on each individual cultivar was composited and used in soil microcosm assays to investigate if the urines derived from each cultivar differentially suppressed nitrification rates. The metabolomes of the composited leaf biomass were also assessed.

### 2.1 Trial details

Grazing trials on 2500 m^2^ plots, were conducted in both autumn (March 2018) and spring (September 2018) at the Agricom Kimihia Research Centre in Canterbury, New Zealand. Each plot was planted with one of nine different plantain cultivars, consisting of lines that varied in geographic origin, flowering date and winter growth activity (Table 1). Plots contained at least 90% plantain at the start of each trial, with the balance being weed grasses. Fertiliser (DAP) was applied prior to sowing across the whole study area at recommended agronomic rates (200 kg ha^-1^). The plots were sown in spring (2017) and were grazed once after establishment when the plants were approximately 3 months old. The autumn grazing trial occurred in March 2018, followed two grazing off periods to even up sward height and then the spiring grazing trial occurred in September 2018. Topping was not required.

**Table 1:** Origin, flowering date and winter growth activity of the plantain cultivars used in the grazing trials.

| Cultivar | Cultivar Origin | Flowering | Winter Activity |
| --- | --- | --- | --- |
| A | New Zealand/Europe | Mid/Late | High |
| B | New Zealand (Mediterranean) | Early | High |
| C | New Zealand (Mediterranean) | Mid | Low |
| D | Northern European | Late | Low |
| E | New Zealand | Early | High |
| F | Unknown | Early | High |
| G | New Zealand | Early | High |
| H | Northern European | Late | Low |
| I | New Zealand | Early | Medium |

In each trial, Romney cross ewes that had previously been grazing ryegrass/clover pasture were transferred onto the plots for a period of 14 days at a stocking rate of five sheep per 2500 m^2^ plot area (Figure 2). Animals were randomised for each trial. After 14 days, spot samples of urine were collected manually from sheep in a sheep handler. For comparison, urine was also collected from five ‘control’ sheep grazing ryegrass-dominated feed; in the autumn this urine was sourced from sheep participating in an indoor feeding trial that were fed a diet of 33% plantain (Cultivar G) and 67% ryegrass (*Lolium perenne* ‘Samson’), while in the spring this urine was sourced from sheep grazing an adjacent plot of 100% ryegrass. Samples of above-ground plant material from each plot were collected at the same time the urine was collected and frozen at −80°C pending analysis. Plant material was ‘plucked’ from the grazing zone from each plot in a transect running diagonally from corner to corner. There were 25 ‘plucks’ per plot to gain one composite sample of leaf biomass per cultivar (plot) per season.

**Figure 2.**
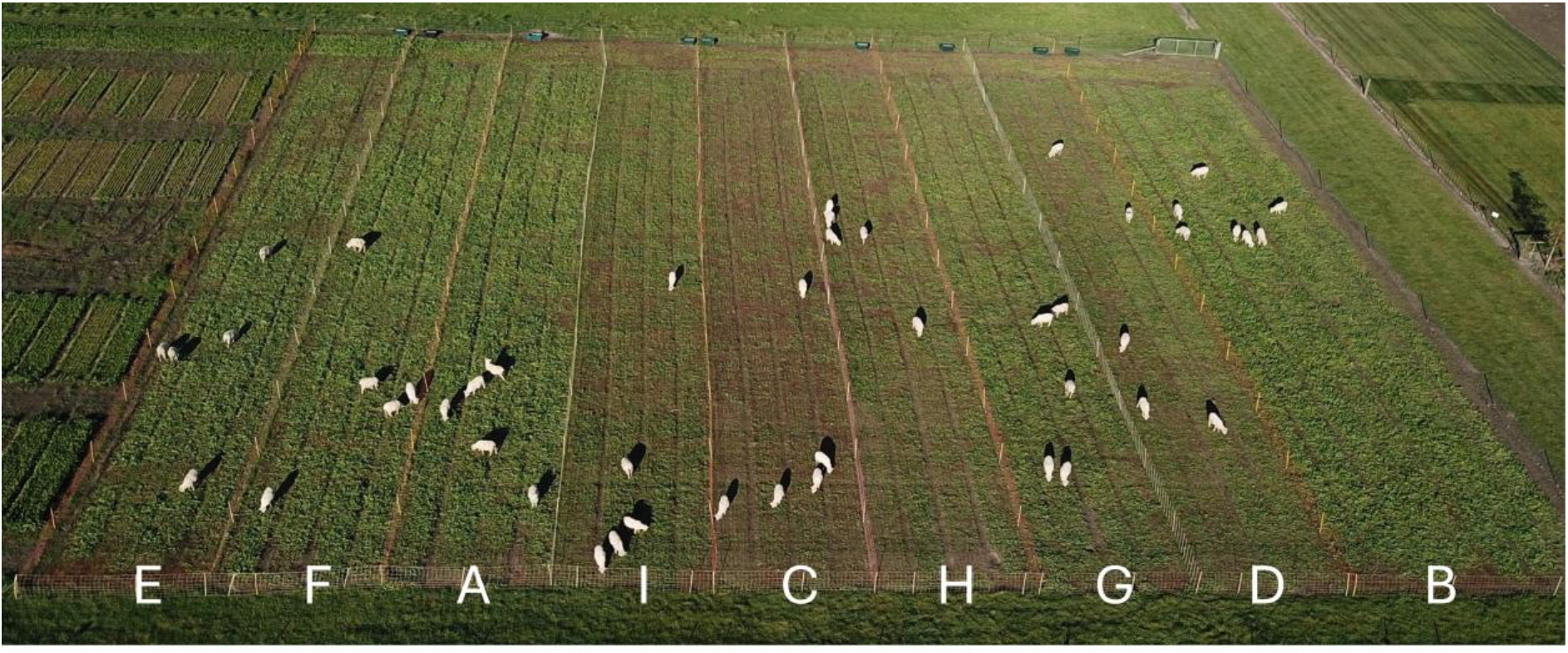
Sheep grazing on plots of different *Plantago lanceolata* cultivars in preparation for urine collection (autumn). This aerial photo was taken towards the end of the grazing period and illustrates the difference in cover provided by each cultivar. Photograph provided by Agricom.

### 2.2 Urine N and urea analyses

Immediately following collection, urine samples were chilled on ice and transported to the laboratory. The urine from each animal was analysed for total N on a QuikChem 8500 Series 2 Flow Injection Analysis System (Lachat Instruments, Loveland, CO, USA) with concentrations of ammonium (NH_4_^+^-N) and nitrate (NO_3_^-^-N) in each sample determined concurrently. The concentration of urea in each sample was determined separately by the method of Talke and Schubert (1965).

### 2.3 Bioassay of the urinary organic fraction (UOF) minus urea

Nitrification inhibition by reconstituted UOF of each of the autumn urines from each individual animal was assessed using a bioassay in which the Griess reaction was used to detect NO_2_^-^ produced by the nitrifying bacterium *Nitrosospira multiformis* (ATCC® 25198™) as it metabolises NH_4_^+^ (O’Sullivan et al., 2017). Inorganic salts and urea were removed from 30 mL urine sub-samples by washing whole urine through a 20 mL Phenomenex StrataX C18 SPE column (Torrance, CA, USA) with three column volumes of Type 1 water (Purite Select Fusion 40, Suez Water, Thame, UK). The adsorbed organic fraction was eluted with 90:10 v/v methanol:water (30 mL), after which the liquid was removed by Speedvac-assisted evaporation (Labconco, Kansas City, KS, USA) and the remaining fraction, labelled as the UOF was retained for use in the bioassay.

To determine an appropriate target concentration for comparing UOF nitrification inhibition rates in the bioassay, a dose response curve was prepared using the UOF derived from animals feeding on Cultivar G. Root exudates from this cultivar exhibited moderate to high BNI in prior hydroponic experiments (data not shown), moderate reduction in NO_3_^-^-N production in soil incubations in both autumn and spring (this work), had good winter activity from an agronomic perspective (as outlined in Table 1), and is a key cultivar within New Zealand plantain breeding programmes. The UOF was tested in multiple assays at concentrations between 0 and 2 mg mL^-1^. This experiment was also used to establish if there was a positive linear relationship between concentration of urinary metabolites and BNI activity, and to determine the maximum inhibitory concentrations at which higher concentrations would lead to static inhibition. Based on these results, and to ensure static inhibition would not be reached, even for high BNI cultivars, UOF fractions from all plantain cultivars and ryegrass were resuspended in water targeting a final concentration of UOF in the bioassay of 0.5 mg mL^-1^.

In a 96-well plate, 35 µL of the *N. multiformis* cell culture (washed and resuspended to an OD_600_ of 0.08) was assayed with a 40 µL aliquot of UOF, with 165 µL *of N. multiformis* growth media added to bring the total well volume to 240 µL. Blank wells contained an aliquot of UOF without cells. An inhibition control of allylthiourea (ATU), a synthetic nitrification inhibitor, was included at a concentration of 10 µM (determined, as the minimum inhibitory concentration for this strain of *N. multiformis*). Assays were performed in triplicate and the production of NO_2_^-^ was followed spectrophotometrically over a 100 min period at 28°C in a temperature-controlled CLARIOstar microplate reader (BMG LABTECH GmbH, Ortenberg, Germany). At 20 min intervals, 10 µL of Griess colour reagent (sulfanilamide 16.2 g L^−1^, N-1-naphthylethylenediamine dihydrochloride 0.8 g L^−1^, concentrated phosphoric acid 41 mL L^−1^) were added to a set of wells and the colour change reaction allowed to proceed for 5 min before the absorbance was read at 548 nm. Absorbance values were converted to NO_2_^-^ concentrations using a standard curve (0 to 2.5 mg L^−1^ NO_2_^−^-N) and nitrification rates were calculated from a linear regression of NO_2_^-^ formation over time. The BNI potential of the UOF was calculated from the decrease in nitrification rate relative to an uninhibited control.

### 2.4 Soil incubation testing nitrification of urinary nitrogen

After appropriate volume subsamples of urine from each sheep were taken for N-chemistry analysis and for the bioassay and metabolomics, the remaining urine from the five sheep from each plot was combined into one bulk sample to provide sufficient volume for a soil incubation study. The bulk urine samples were re-analysed for nitrogen using the methods described above. To ensure that urinary N inputs were standardised across treatments, the concentration of N in each urine sample was adjusted to that of the lowest total-N (bulk) urine by addition of deionised water before use in the incubation study. Bulk urine N chemistry and dilution information can be found in Supplementary Table 1.

A Templeton silt loam soil (Udic Haplustept, USDA) from a site under low-input, long-term ungrazed Italian ryegrass (*Lolium multiflorum* ‘Andy’) was used for the incubation experiments. The soil was sampled from the New Zealand Bioeconomy Science Institute farm in Lincoln, Canterbury, New Zealand (43°38’17.8”S 172°28’27.4”E). Multiple soil cores were taken to a depth of 5 cm, mixed thoroughly into one composite sample, passed through a 4 mm sieve and air-dried. This soil was a slightly acidic silt-loam of pH 5.6 (1:2.5 soil water ratio, Devey et al. (2010)) with a carbon content of 29 g kg^-1^ (combustion method using a LECO instrument, Skjemstad and Baldock (2008)), a low background mineral N content of 1.7 mg N kg^-1^ (flow injection analysis method using a Lachat Quikchem 8500, Keeney and Nelson (1982)) and a nitrification potential of 9 mg kg^-1^ day^-1^ (measured using a potential nitrification rate – PNR - assay, Drury et al. (2008)). Sub-samples of soil (20 g air-dry) were weighed into plastic containers and 6.3 mL of urine (10 treatments) or water (no urine control) was added dropwise using an electronic pipette in titrate mode. Five experimental replicates of each treatment and the control treatment were destructively sampled at 1, 7, 14, 21 and 28 days (a total of 275 microcosms). Urine from sheep in the autumn grazing trial was added to microcosm soil at a concentration of 310 mg N kg^-1^, while the spring urine was added at 362 mg N kg^-1^. These N additions were defined by the N concentration within lowest-N (bulk) urine and the 6.3 mL volume of liquid calculated to bring the soil to field capacity (defined as the soil water content at −10 kPa; actual water content of 32%). The microcosms were covered with Parafilm®, which was pierced to allow for gas diffusion. Microcosms were placed in 5.5 L boxes where each box contained 11 soil microcosms (urine treatments plus control – sub-plots) and represented a timepoint (main plots). Boxes were split across 5 incubator shelves (blocks), with each timepoint and associated treatments represented on each shelf. The incubator was set at 20°Cand the microcosms soil was maintained at −10 kPa during the incubation by the addition of water as required. At each time point, a box was removed from each shelf, and 5 g soil was extracted from each microcosm with 25 mL of 2 M KCl and the filtered extract analysed for NO_3_^-^-N and NH_4_^+^-N on a QuikChem 8500 Series 2 Flow Injection Analyser (Lachat Instruments, Loveland, CO, USA).

Statistical tests on the incubation data were performed using Genstat 64-bit Release 19.1 (VSN International Ltd, Hemel Hempstead, UK). Differences between treatment means at each timepoint were tested by one-way analysis of variance (ANOVA), and if treatment effects were significant at P < 0.05 and after Bonferroni correction, least significant difference (LSD) was used to separate means. LSD was not intended to be used to compare the plantain cultivars against each other, rather it compared the nine cultivars versus the no-urine control or the ryegrass urine treatment. Any trends that suggested significant differences among plantain cultivars were referred for further analysis in another research and breeding programme.

### 2.5 Plant material metabolite extraction

Frozen plant material from both trials was freeze-dried then crushed, thoroughly mixed and ∼10 mg weighed into 2 mL polypropylene tubes. One mL of 96% ethanol in water was added, the samples vortexed briefly then left for 16 h on an orbital shaker. The samples then were placed in floating tube holders and sonicated for 20 min at 37 Hz in a water filled ultrasonic bath before centrifugation at 15700 × g for 5 min. A 200 μL aliquot of supernatant was added to a filter vial (Thompson™ Single Step®, 0.22 mm PVDF, low evaporation crimp top) along with the addition of 200 μL of Type 1 water. The filter unit was pushed in part way to just seal then the sample was briefly vortexed to mix prior to completing filtration.

### 2.6 Urine metabolite extraction

Urine subsamples from both trials were set aside for mass spectrometry-based untargeted metabolomic analysis. Samples were chilled at 2°C for a maximum of 24 h before a 200 μL aliquot of urine was added to a filter vial followed by the addition of 200 μL of cold acetonitrile. The filter unit was pushed in part way to just seal then the sample was briefly vortexed to mix prior to completing filtration. Where urine volume from individual sheep was larger, additional subsamples were collected and included as technical replicates in the analysis.

### 2.7 Plant and urine metabolomics

Ultra-high-performance LC-MS analysis was performed on a Q Exactive™ Plus Orbitrap (HR/AM) LC-MS/MS coupled with a Vanquish™ UHPLC system (Binary Pump H, Split Sampler HT, Dual Oven) that was calibrated immediately prior to sample analysis batch with Pierce™ LTQ ESI positive and negative ion calibration solutions (ThermoScientific, CA, USA). Samples were analysed by four HRAM LC-MS methods; reverse phase (C) and aqueous phase (H) with negative (n) and positive (p) ionisation modes, generating four datasets (Cp, Cn, Hn, Hp) to cover a large range of metabolite attributes. An aliquot (2 μL) of each sample and appropriate reference materials (aucubin, catalpol and verbascoside in the plant material) was analysed. Reverse phase chromatography (C) used an Accucore Vanquish C18 column (1.5 µm, 100 mm × 2.1 mm, ThermoScientific) maintained at 40 °C, with a flow rate of 400 µL/min and a mobile phase consisting of 0.1 % formic acid in type 1 water (A) and 0.1 % formic acid in acetonitrile (B). The following gradient was applied: 0–1 min/0%B, linear increase to 7 min/50%B, linear increase to 8 min/98%B, isocratic to 11 min/98%B, equilibration 11– 12min/0%B, isocratic to end 17 min/0%B. Normal phase chromatography (H) used a Hypersil Gold HILIC column (1.9 µm, 100 mm × 2.1 mm, ThermoScientific), maintained at 55°C, with a flow rate of 400 µL/min and a mobile phase consisting of 0.1 % formic acid in acetonitrile (A) and 5 mM ammonium acetate in water (B). The following gradient was applied: 0–1 min/5%B, linear increase to 12 min/98%B, isocratic 16min/98%B, equilibration 16–17 min/5%B, isocratic to end 20min/5%B. The eluents from (H) and (C) chromatography were scanned from 0.5–16 min and 0.4–11.5 min, respectively, by API-MS (Orbitrap) with heated electrospray ionisation (HESI) at 350°C in the negative and positive mode with capillary temperature of 320°C. Data were acquired for precursor masses from m/z 80–1200 amu (H) and m/z 100–1500 (C) at 70K resolution (AGC target 3e^6^, maximum IT 100ms, profile mode) with data dependent ms/ms for product ions generated by normalised collision energy (NCE:35, 45, 65) at 17.5K resolution (TopN 10, AGC target 2e5, Maximum IT 50 ms, Isolation 1.4 *m/z*).

Reference materials verbascoside, aucubine, catalpol (CAS; 61276-17-3; 479-98-1; 2415-24-9) were sourced from Phyproof™ (catalogue numbers; 89289; 89163; 89595, respectively). Data were processed using Thermo-Fisher Compound Discoverer (version 3.3) software. Plant compound-related, mass-feature datasets were filtered and processed via differential analysis with correlative discrimination against ovine urine data and soil assay results at 21 and 28 days (NO_3_^-^-N, 92 to 288 mg kg^-1^) for each of the plantain cultivars. For plantain biomass metabolites, the area response of putatively identified major leaf compounds after differential analysis was normalised relative to the highest concentration of verbascoside measured to give crude estimates of concentrations allowing comparison across all cultivars. These crude estimates were converted to percentages with heat maps and principal components analysis (PCA) used to visually compare seasonal and cultivar differences. For urine metabolites, differential analysis and discrimination included data from the autumn UOF bioassay used to rank BNI potential, and area response ratios were used to normalise data. It should be noted that the use of autumn UOF bioassay results for discriminating spring urines probably reduced the number of correlating metabolites because the BNI potential and nitrification profiles for each cultivar was not consistent between seasons. PCA of metabolite chemistry was performed using PRIMER 7 with PERMANOVA add-on (Primer-E Ltd) with square root data transformation, Euclidean distance matrices and built-in default analysis parameters.

## 3 Results

### 3.1 Urine N analysis and nitrification in urine-treated soil microcosms

The urine N analyses for groups of sheep fed one of nine different cultivars of plantain in grazing trials conducted in autumn and spring are summarised in Table 2. Total N in the autumn ranged from a low of 1061 mg L^-1^ in urine derived from Cultivar G to a high of 4932 mg L^-1^ in the ryegrass urine. Urinary N range was much higher in the spring with total N in the urine from sheep fed Cultivar C the lowest (1273 mg L^-1^) while the ryegrass urine was again the highest (9851 mg N L^-1^). A higher proportion of the total N was represented in the non-urea nitrogenous compounds (NUNC, calculated as the difference between total N and urea-N – Table 2) fraction in the spring. There was a strong positive correlation (R^2^ >0.99) between total N and urea concentration, but no other obvious relationships were observed between any of the N pools.

**Table 2:** Mean and standard deviation (SD) for urine-N parameters in autumn and spring measured from sheep after 14 days of grazing. Urine was collected from sheep eating one of nine different plantain cultivars with a stocking rate of five sheep per 2500m^2^ plot area. Cultivars with medium to high winter activity highlighted in grey. Non-urea nitrogenous compounds (NUNC) predominate within the urinary organic fraction (UOF) and represents Total-N minus Urea-N. The Soil-NO_3_^-^ figures presented is nitrate measured in incubated soil microcosms sampled destructively at day 28. For comparison, urine was also collected from five ‘control’ sheep grazing ryegrass-dominated feed; in the autumn this urine was sourced from sheep participating in an indoor feeding trial that were fed a diet of 33% plantain (Cultivar G) and 67% ryegrass (*Lolium perenne* ‘Samson’), while in the spring this urine was sourced from sheep grazing an adjacent plot of 100% ryegrass.

| Cultivar | Autumn |  |  |  |  |  |  |  | Spring |  |  |  |  |  |  |  |
| --- | --- | --- | --- | --- | --- | --- | --- | --- | --- | --- | --- | --- | --- | --- | --- | --- |
|  | Total N (mg N L<sup>-1</sup>) |  | NH<sub>4</sub><sup>+</sup>-N (mg N L<sup>-1</sup>) |  | Urea-N (mg N L<sup>-1</sup>) |  | NUNC<br>(mg N L<sup>-1</sup>) | Soil-NO<sub>3</sub><sup>-</sup><br>(mg kg<sup>-1</sup>) | Total N (mg N L<sup>-1</sup>) |  | NH<sub>4</sub><sup>+</sup>-N (mg N L<sup>-1</sup>) |  | Urea-N (mg N L<sup>-1</sup>) |  | NUNC<br>(mg N L<sup>-1</sup>) | Soil-NO<sub>3</sub><sup>-</sup><br>(mg kg<sup>-1</sup>) |
|  | Mean | SD | Mean | SD | Mean | SD |  |  | Mean | SD | Mean | SD | Mean | SD |  |  |
| A | 1833 | 240 | 47 | 3 | 1694 | 175 | 139 | 275 | 4385 | 949 | 62 | 53 | 3768 | 1221 | 617 | 232 |
| B | 1336 | 763 | 36 | 20 | 1127 | 627 | 209 | 261 | 5807 | 1384 | 55 | 12 | 4494 | 1283 | 1312 | 250 |
| C | 2848 | 454 | 74 | 17 | 2610 | 614 | 258 | 238 | 1273 | 362 | 56 | 41 | 924 | 202 | 349 | 260 |
| D | 1400 | 697 | 36 | 35 | 1238 | 567 | 162 | 237 | 2102 | 1312 | 146 | 199 | 1751 | 1158 | 351 | 189 |
| E | 1898 | 898 | 56 | 22 | 1606 | 794 | 291 | 248 | 6685 | 1562 | 45 | 14 | 6112 | 1553 | 573 | 266 |
| F | 1250 | 257 | 59 | 32 | 1146 | 305 | 104 | 253 | 4205 | 1548 | 49 | 30 | 3810 | 1513 | 395 | 248 |
| G | 1061 | 221 | 12 | 13 | 776 | 168 | 285 | 247 | 5190 | 327 | 57 | 31 | 4351 | 391 | 839 | 231 |
| H | 3013 | 967 | 71 | 14 | 2873 | 968 | 140 | 231 | 2109 | 1435 | 102 | 73 | 1527 | 1173 | 582 | 215 |
| I | 1850 | 1850 | 114 | 114 | 1761 | 1761 | 89 | 201 | 5456 | 1920 | 61 | 33 | 4976 | 1833 | 480 | 246 |
| Ryegrass | 4932 | 1089 | 831 | 388 | 4757 | 1316 | 201 | 270 | 9851 | 2909 | 91 | 52 | 8882 | 2518 | 969 | 288 |

The results of soil incubations that recorded the production of NO_3_^-^-N through nitrification after application of urine from groups of sheep fed one of nine different plantain cultivars are shown in Figure 3. In the autumn trial (Figure 3a), large plantain cultivar effects on soil nitrification were observed, particularly in the early phase of the incubation. All the urines delayed nitrification in the first 7–10 days compared with the no urine control. This delay was eventually overcome at around 10 days for Cultivars A, B and C and 12 days for the others except Cultivar I, which surpassed the no-urine control at approximately 16 days. This delayed or slowed nitrification was also evident in the ammonium (NH_4_^+^-N) data (Supplementary Figure 1).

**Figure 3:**
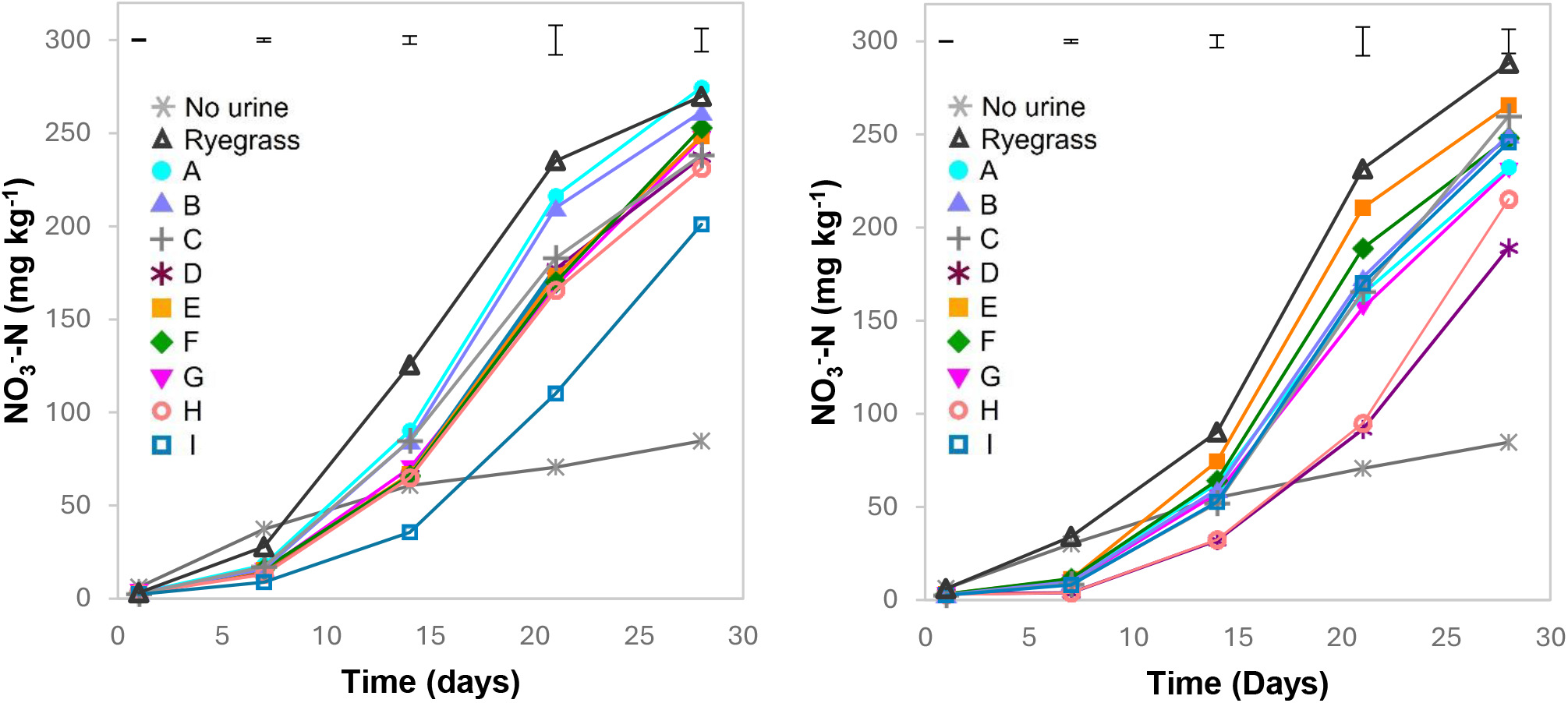
Nitrate produced in soil microcosms over 28 days when urine from sheep in the autumn plantain grazing trial was added at a concentration of 310 mg N kg^-1^ (a), or urine from animals in the spring plantain grazing trial was added at 362 mg N kg^-1^ (b). The autumn ryegrass treatment was sampled from sheep housed indoors and fed a diet consisting of 33% Cultivar G and 67% ryegrass, Cultivar Thirty-three. The spring ryegrass treatment was sampled from sheep fed outdoors on 100% ryegrass, cultivar Thirty-three. Plantain cultivar treatments A-I were fed outdoors with swards containing >90% plantain. The error bars represent LSDs (p<0.05) at each incubation time. Figure 2a is a permitted reproduction of data published in the IGC Proceedings 2023, Peterson et al. (2023).

Nitrate-N produced over the 28-day incubation ranged from 201 to 275 mg kg^-1^ soil (Table 2) with urine from Cultivar I, having the greatest inhibition producing approximately half the NO_3_^-^-N compared with the urines from other cultivars up to day 21. Cultivars D, E, F, G, H and eventually C did not appear to be significantly different from each other and could be considered to have moderate nitrification inhibition within the context of this study. Cultivars A and B had the lowest inhibition, were separated from this moderate inhibition group, but not different from each other. All plantain cultivars were significantly different to the ‘predominantly’ ryegrass urine treatment, which showed the most NO_3_^-^-N production, despite a third of the diet being the moderately inhibitory plantain Cultivar G. Between 21 and 28 days, NO_3_^-^-N production rates slowed for cultivars A, B, C, D and H. Rates of NO_3_^-^-N production for Cultivar I increased to be equivalent to E, F, and G, but overall NO_3_^-^-N produced did not surpass the other cultivars at day 28. Nitrate production for Cultivars A and B was equivalent to ryegrass by day 28.

Similarly, in the spring trial (Figure 3b), large plantain cultivar effects on soil nitrification were observed in the early phase of the incubation. All urines delayed nitrification in the first 7–10 days compared with the no urine added control with this delay eventually overcome by day 14 for all cultivars except E (less delay, 10 days, lower inhibitory capacity), D and H (greater delay, both ∼17 days, highest inhibition capacity). In spring, NH_4_^+^-N was always lower in the soils treated with plantain urines and the nitrification delay was supported until day 14 (Supplementary Figure 1). Nitrate-N produced over the 28-day incubation ranged from 189 to 288 mg kg^-1^ soil (Table 2). Urine from sheep grazing Cultivars D and H yielded results that trended differently from the other cultivars, producing less than half of the NO_3_^-^-N than the next “best” cultivar (G) after 21 days. Between 21 and 28 days, NO_3_^-^-N production rates slowed except for Cultivars D and H where rates increased, but NO_3_^-^-N produced did not surpass that produced from urines derived from the other cultivars.

In the spring a weak negative correlation (R^2^ = −0.57) was observed between the concentration of non-urea nitrogenous compounds (NUNC) in the urine and the amount of NO_3_^-^-N produced in the soil microcosms after 28 days; the amount of NO_3_^-^-N in the soil weakly correlated (R^2^ = 0.49) with total N in the spring urine samples, but no obvious relationship was observed between NO_3_^-^-N in the soil and total N or NUNC in the autumn urine samples.

### 3.2 Bioassay of the autumn urine organic fractions to guide metabolomic analysis

Urine derived from sheep fed Cultivar G was associated with the lower NO_3_^-^-N production rates in the soil incubation assays over 28 days in autumn (and spring). Urine from this cultivar was used to benchmark the concentration of UOF required in the bioassay to enable comparison between the different UOF from the groups of sheep fed one of the nine different plantain cultivars. In this benchmarking ‘dose response’ test, the final concentration of Cultivar G UOF varied between 0 and 2 mg mL^-1^ resulting in a positive correlation between increasing UOF concentration and nitrification inhibition with 100% inhibition reached between 1.25 and 1.5 mg mL^-1^ UOF (Supplementary Figure 2). Based on this result, a final UOF bioassay concentration of 0.5 mg mL^-1^ was chosen to allow comparison of UOF from different urines derived from sheep in the autumn grazing trial.

Bioassay results for the nine urine samples from the autumn grazing trial are presented in Supplementary Figure 3a. Inhibition caused by the ryegrass urine was excluded from the analysis due to excessive colouration in the sample interfering across wavelengths measured within the colorimetric assay. This was caused by the presence of contaminant faecal matter. The bioassay showed a similar order of cultivars with respect to their nitrification inhibition efficacy when compared with the soil incubations. There was a reasonable inverse correlation (R^2^ = 0.75) between % inhibition and NO_3_^-^-N production in the soil microcosms after 28 days (Supplementary Figure 3b). Bioassay results were then carried forward as part of the correlative discriminant analysis of urinary metabolites from the spring grazing trial.

### 3.3 Metabolomic analysis of leaf extracts from autumn and spring

A total of 1461 mass features were recorded in the leaf material across the plantain cultivars. After filtering, 497 mass features were considered for statistical correlation against NO_3_^-^ -N produced in soil incubations. Of these, only 14 correlating mass features were shared by all cultivars, and these were then putatively identified using a plantain compound database constructed from literature reports of these compounds (SciFinder™), manual interpretation of structural fragments and comparison with those identified in other Plantaginaceae (Table 3).

**Table 3.** Major discriminating compounds identified in the leaves of all plantain cultivars in autumn and spring grazing trials using reference standards^†^ and putatively interpreted from liquid chromatography combined with mass spectrometry (LC-MS) spectral data. Unless indicated via references dated after the year 2000, all previous identifications of these compounds are from a comprehensive study of the chemotaxonomy of *Plantago* species by Rønsted et al. (2000) and references therein. CAS = Chemical Abstracts Service.

| ID | Name | CAS # | Class* | Previous identifications |
| --- | --- | --- | --- | --- |
| L1 | Campneoside I | 95519-12-3 | PhG | *P. asiatica*; *P. depressa* (Qi et al., 2012) |
| L2 | Plantamajoside | 104777-68-6 | PhG | *P. arborescens*; *P. asiatica*; *P. lanceolata*; *P. major*; *P. media* (Andary et al., 1988); *P. ovata*; *P. webbii* |
| L3 | (R)-β-Hydroxyverbascoside | 91050-24-7 | PhG | *P. lanceolata* (Fleer and Verspohl, 2007) |
| L4 | Oxoverbascoside | 149507-92-6 | PhG | *P. depressa* (Nishibe et al., 1993) |
| L5 | Verbascoside<sup>†</sup> | 61276-17-3 | PhG | All Plantaginaceae (Andary et al., 1988) |
| L6 | Desacetylhookerioside | 180028-17-5 | IrG | *P. altissima* (Jensen et al., 1996); *P. subulata* |
| L7 | Globularin | 1399-49-1 | IrG | *P. altissima* (Handjieva et al., 1991); *P. lanceolata* (Handjieva et al., 1991) |
| L8 | Dumuloside | 613246-43-8 | IrG | Reported within Plantaginaceae (Kirmizibekmez et al., 2003) |
| L9 | Asperuloside | 14259-45-1 | IrG | *P. altissima*; *P. bellardi*; *P. lanceolata* (Bianco et al., 1984); *P. major* |
| L10 | Mussaenosidic acid | 82451-22-7 | IrG | *P. alpina* (Jensen et al., 1996); *P. arborescens*; *P. nivalis*; *P. sempervirens*; *P. webbii* |
| L11 | Geniposidic acid | 27741-01-1 | IrG | *P. alpina* (Jensen et al., 1996); *P. asiatica* (Endo et al., 1981); *P. bellardi*; *P. lundborgii*; *P. reniformis*; *P. major* |
| L12 | Arborescosidic acid | 325798-51-4 | IrG | *P. atrata*; *P. maritima*; *P. stauntonii*; *P. subulata* |
| L13 | Catalpol<sup>†</sup> | 2415-24-9 | IrG | Most Plantaginaceae |
| L14 | Aucubin<sup>†</sup> | 479-98-1 | IrG | Most Plantaginaceae |

\* PhG – phenylethanoid glycoside; IrG – iridoid glycoside
<sup>†</sup> Identified against reference standards

The known bioactive plantain compounds aucubin, catalpol and verbascoside were quantified from leaf material sampled in both autumn and spring as mg g^-1^ dry matter and are presented in Figure 4. For each cultivar, verbascoside concentrations were between 2.9 and 17.7 times higher in spring compared to autumn while aucubin was between 1.1 and 3.8 times higher in spring versus autumn. Catalpol was up to 7.5 times higher in spring compared to autumn for Cultivar G, while Cultivar A had 0.8 mg g^-1^ in spring and 0 mg g^-1^ in autumn. Visual trends suggested a tenuous inverse relationship between the concentration of NO_3_^-^-N accumulated after 28 days in the urine-treated soil incubations and the concentration of verbascoside in the leaf in spring, but this could not be statistically tested. Aucubin was present at concentrations up to 10 mg g^-1^ irrespective of season or cultivar except for Cultivar I in spring. Ignoring Cultivars I and C in spring, catalpol also had a tenuous inverse visual trend against concentration of NO_3_^-^-N accumulated in soil after 28 days. Again, this trend could not be statistically tested.

**Figure 4:**
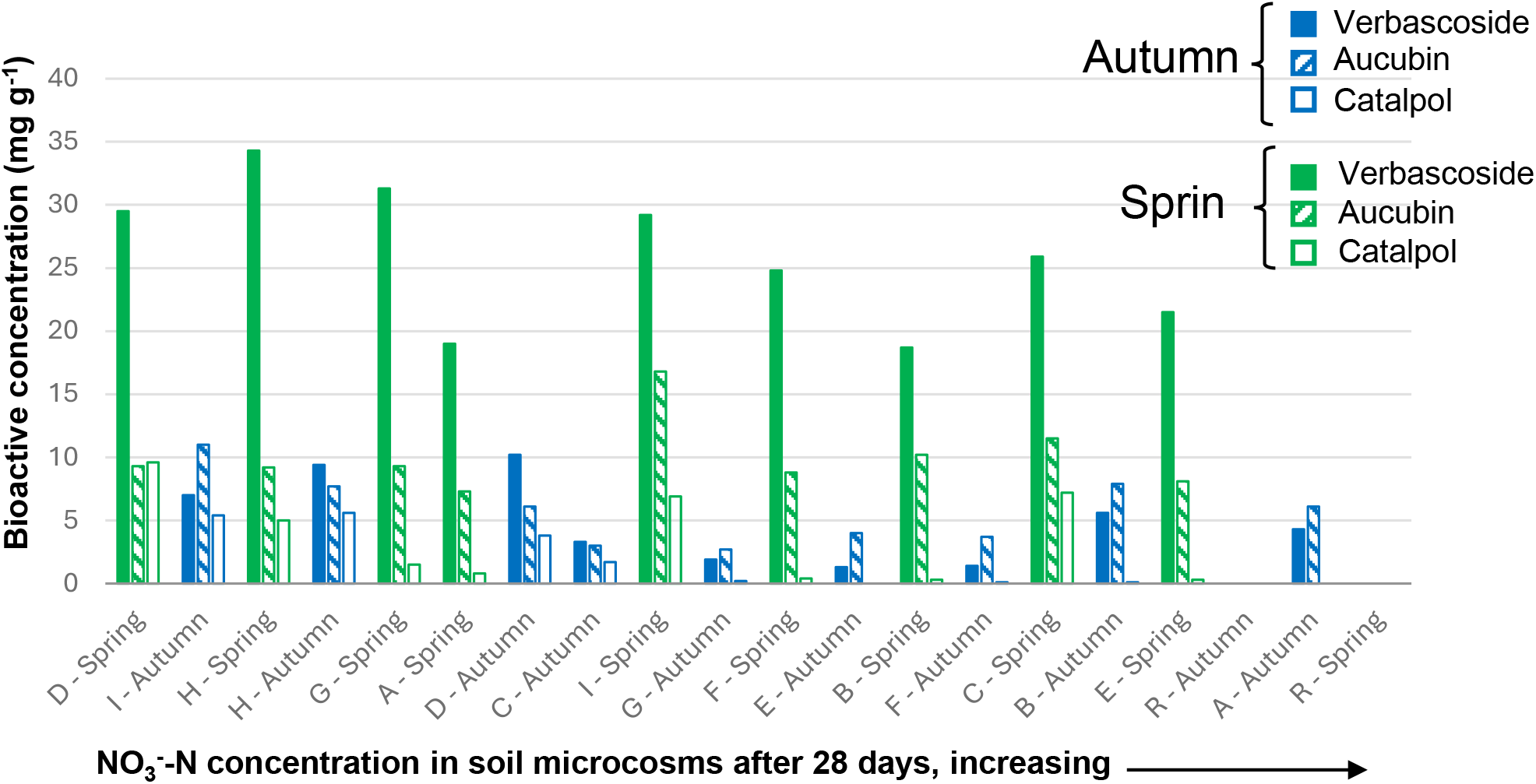
Concentrations of three key plantain bioactive compounds extracted from leaf biomass for Cultivars A–I in spring (green bars) and autumn (blue bars). Verbascoside (filled bars), aucubin (hatched bars) and catalpol (open bars). The position on the x-axis represents cultivar ranking in terms of nitrate concentration in the soil microcosms, lowest to highest (left to right) after 28 days of incubation. R = Ryegrass.

To enable inferences to be made about their influence on BNI, the abundance (expressed as the area under the curve) of each leaf metabolite putatively identified in Table 3 was normalised relative to the highest concentration of verbascoside observed across the two grazing trials (35 mg g^-1^ dry matter in Cultivar H when grazed in the spring, Figure 4). This provided a normalised area response in the absence of quantification against standards. A heat map representing these normalised area response concentrations as percentages is presented in Figure 5. Leaf concentrations of all metabolites are generally higher in spring with Cultivar G having several standout metabolites (e.g. plantamajoside - L2 and globularin – L7) that are different than the other cultivars. Urines from cultivars with the lowest nitrate production when applied to soil - Cultivar I in autumn, and D and H in spring - are derived from plants that had the highest leaf aucubin in autumn and verbascoside in spring.

**Figure 5:**
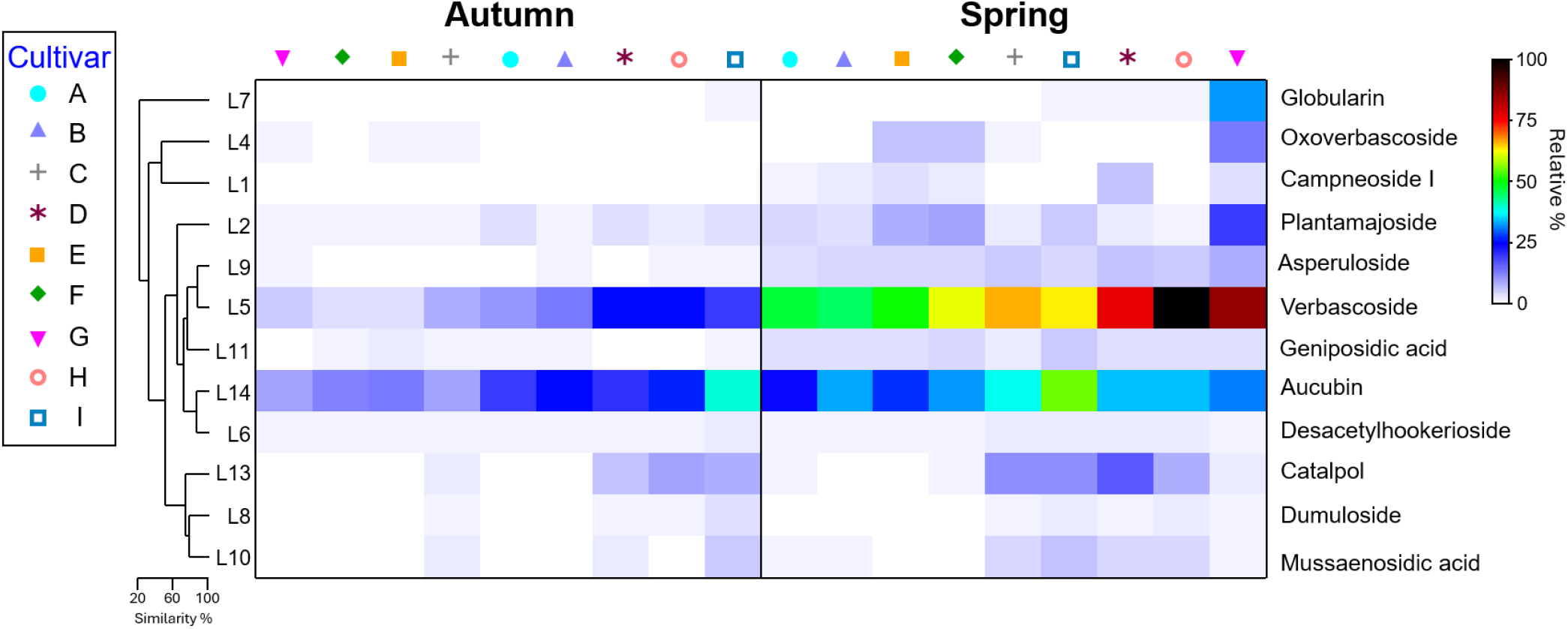
Heat map of the seasonal abundance of leaf metabolites L1 to L14 relative to verbascoside for Cultivar H where 100% = 34.3 mg g^-1^. Clustering dendrogram Similarity % represents how closely grouped the metabolites are between cultivars in each season and between seasons. L2 and L3 were combined, as they have similar structures aside from the sugar group, while L11 and L12 were combined as they are structural isomers.

In addition to the heat map, an exploratory principal components plot suggested that alongside verbascoside (L5 – as observed in Supplementary Figure 4), the phenylethanoid glycosides (PhGs), plantamajoside (L2), (R)-ß-hydroxyverbascoside (L3) and oxoverbascoside (L4) may be associated with reduced NO_3_^-^-N concentrations in the soil incubations observed between seasons specifically for cultivar G, but also A, B, and F with a lesser role for campneoside I (L1). The greater abundance of these PhGs in the spring appears to be related to the enhanced BNI effect of the urine from the sheep fed these cultivars on soil NO_3_^-^-N production, with smaller contributions from the iridoid glycosides (IrGs), globularin (L7), asperuloside (L9), geniposidic acid (L11) and arborescosidic acid (L12) (Figure 5, Supplementary Figure 4).

Based on vector direction and length, lower NO_3_^-^-N between autumn and spring for Cultivars D and H, the general decrease in NO_3_^-^-N observed across all cultivars in autumn, and the marked difference between Cultivar I and other cultivars in autumn and spring, may be linked to IrGs desacetylhookerioside (L6), dumiloside (L8), mussaenosidic acid (L10), catalpol (L13) and aucubin (L14) (Supplementary Figure 4).

### 3.4 Metabolomic analysis of spring urine

The major metabolites in the urine found to be associated with changes in the amount of NO_3_^-^ -N produced in soil incubations due to the consumption of plantain are presented in Table 4. The dataset presented here represents one of four chromatography protocols run with reverse-phase data based on negative ions best representing the metabolites of interest. A total of 16,651 mass features were recorded. After filtering data for mass feature significance (Adj.P-value<0.005, ratio >10, area >0.1% max area feature) and correlating peak area with ‘soil NO_3_^-^-N’ values and bioassay results for each cultivar, 244 mass features were retained. Nineteen correlating urinary metabolites common to all urines (with respect to soil NO_3_^-^-N concentrations and bioassay results) were then putatively identified based on spectral data considered in reference to possible derivation from the main leaf compounds expressed in plantain (verbascoside, aucubin and catalpol) and to those known to be urine metabolites (http://lmdb.ca/metabolites - Goldansaz, S.A. et al., 2017). Metabolites could be separated as either urine metabolites, aromatic metabolites, or phenylethanoid (PhG) metabolites.

**Table 4.**
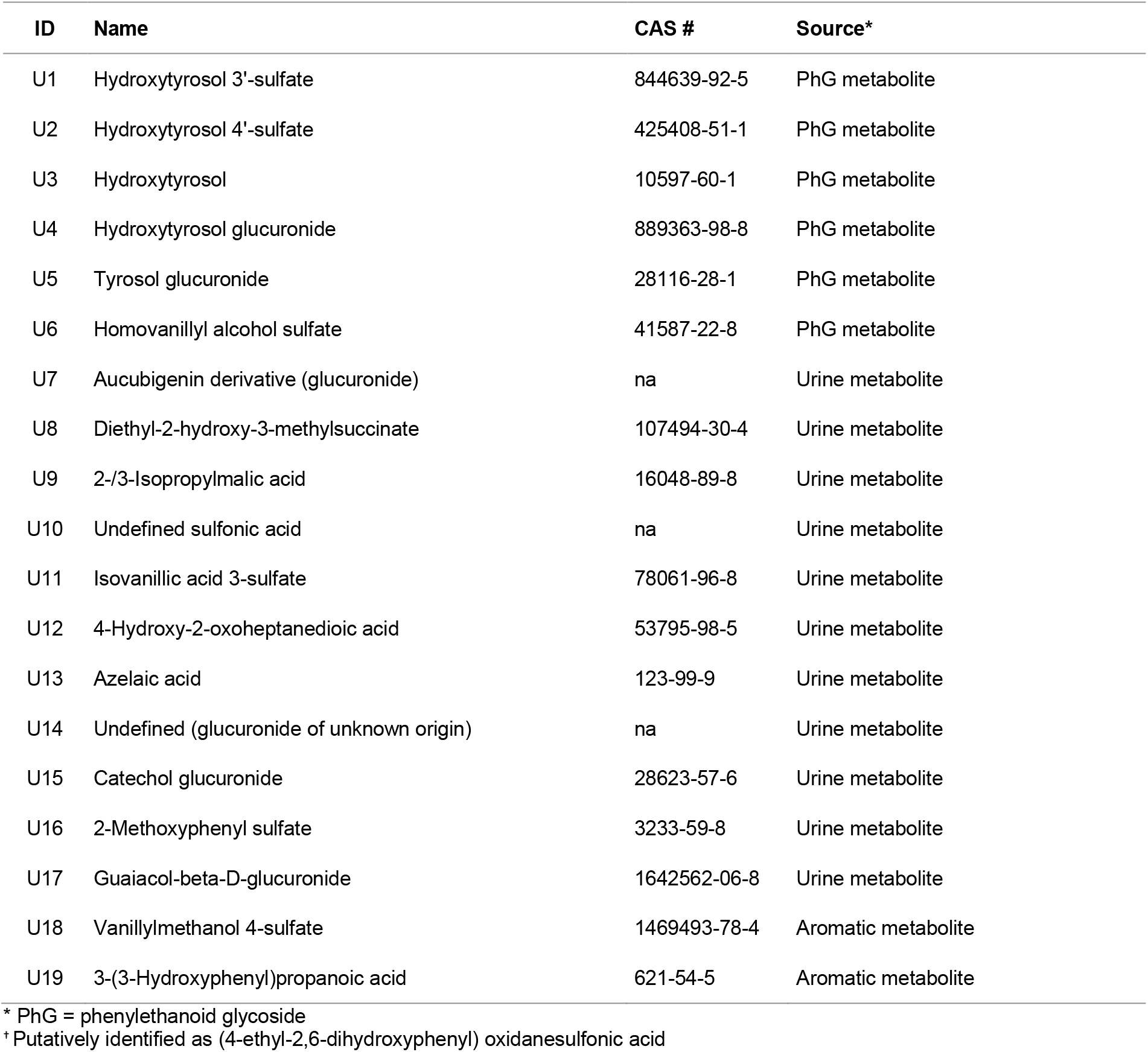
Major discriminating metabolites identified across all urines from sheep fed plantain in both the autumn and spring grazing trials. Compounds were putatively identified based on spectral data considered in reference to the main leaf compounds expressed in plantain (verbascoside, aucubin and catalpol). CAS = Chemical Abstracts Service, na = not available.

PCA of percentage peak area ratios of urinary metabolites from animals in the spring grazing trial indicated substantially different profiles for urine derived from animals grazing Cultivars D, H and I, with D and H having the lowest production of NO_3_^-^-N across all cultivars for both seasons, while Cultivar I exhibited the lowest NO_3_^-^-N production in autumn (Figure 6 and Table 2). For Cultivar H, and to some extent I, the key metabolites associated with their distinction from other urine secondary metabolite profiles were the PhG derivatives: hydroxytyrosol 3’-sulfate (U1), hydroxytyrosol 4’-sulfate (U2), hydroxytyrosol glucuronide (U4), and to lesser extent the aucubigenin derivative (U7). Compounds 4-methoxy-3-(sulfooxy) benzoic acid (U11) and vanillylmethanol 4-sulfate (U18) may also be important as they have similar vector length and direction to U1, U2, and U4. For Cultivar D, which exhibited the lowest NO_3_^-^-N production in the entire study, and again to some extent Cultivar I, the key urinary metabolites were catechol glucuronide (U15), 2-methoxyphenyl sulfate (U16) and guaiacol-ß-D-glucuronide (U17).

**Figure 6:**
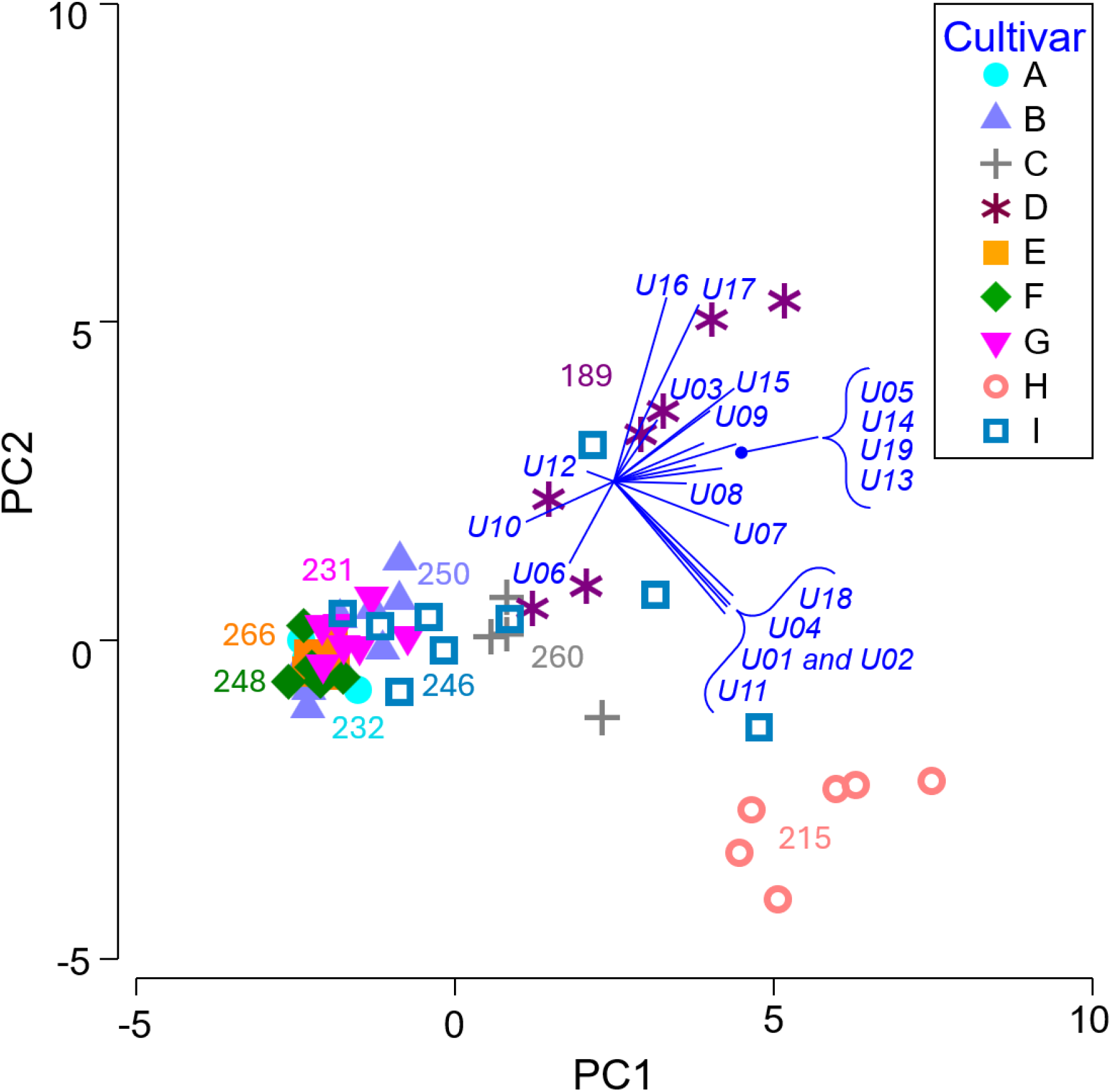
Principal components analysis (PCA) of percentage peak area ratios of metabolites U1 to U19 presented in Table 4 identified in the urine of sheep fed plantain Cultivars A–I in the spring grazing trial, with technical replication where urine volume collected allowed. 42.6% of variation is captured by the PC1 axis and 14.9% by PC2. The PC3 axis captured a further 9.7%. Numbers associated with each set of cultivar data points are the concentrations of nitrate (mg kg^-1^) measured in the soil microcosms after 28 days of incubation with spring urines from groups of animals fed one of the nine different plantain cultivars. The blue vector fan indicates increasing percentage peak area ratios of each metabolite from the centre out, with length and the direction being an indicator of ‘influence’ that each metabolite has with respect to where data points are positioned in the plot relative to each other.

The only other metabolites identified that were shared by Cultivars A and G with relatively low NO_3_^-^-N production were hydroxytyrosol (U3) and 4-hydroxy-2-oxoheptanedioic acid (U12), but both metabolites had little influence within the PCA. Figure 6 represents a projection of the PC1 and PC2 axes which capture 57.5% of the variation, but the PC3 axis captured a further 9.7%, better resolving differences between cultivars C, D, H and I - partially through metabolites U8, U14, but more specifically through the aucubigenin derivative (U7). Hydroxytyrosol (U3) was also moderately high in urine derived from Cultivars B and C, that presented good nitrification inhibition in the bioassay, although it should be noted that the bioassay only tested UOF from animals in the autumn grazing trial (see Supplementary Figures 2 and 3).

## 4 Discussion

### 4.1 Biological nitrification inhibition (BNI) in urine-treated soil microcosms

Groups of sheep (n=5) were fed one of nine different plantain cultivars with the expectation that the resulting urine would chemically differ and that these differences would contribute to differences in BNI potential. The nitrogen concentrations in the sheep urine are close to, or within the ranges reported by other authors such as 1.4–17.8 g N L^-1^ in urine of sheep grazing a ryegrass/cocksfoot/white clover pasture (Hoogendoorn et al., 2010) or 3.0–13.7 g N L^-1^ for sheep on a ryegrass/white clover diet (Bristow et al., 1992). The difference in total N for the cultivars when fed in autumn compared with spring was large and tended to follow that if total N was low in autumn, then it was high in spring and vice versa (see Table 2). Interestingly, this vice versa seasonal difference in N tended to coincide with whether a variety is winter active or not. This will likely affect metabolite expression dynamics within the plant, feed composition and urine chemistry at the time of sampling, which will have a bearing on nitrification inhibition capacity. Nitrate produced from urine from animals fed 100% ryegrass was higher than all the plantain cultivars, consistent with prior studies (Judson et al., 2019; Peterson et al., 2022; Peterson et al., 2023) reviewed in Pinxterhuis et al. (2024) and Eme & Roche (2025).

The delays in nitrification observed after urine was added to soil microcosms were at least 3 days greater than those observed by Peterson et al. (2022). This was using the same soil type, sampled from the same location, with the same incubation methodology and a similar grazing period prior to urine collection. Within the current study, longer nitrification delays would have been influenced by lower initial concentrations of NH_4_^+^-N and urea-N in the animal urine, which also resulted in lower final concentrations of NO_3_^-^-N in the soil. In both autumn and spring, the original concentration of a NH_4_^+^-N in bulk plantain urines is lower than in ryegrass (see Supplementary Table 1). Once the urine is applied to soil it appears that urea in ryegrass urine is rapidly ammonified, coupled with higher initial nitrification rates which is supported by nitrate production data (see Figure 2). In autumn, aside from Cultivars A and B, NH_4_^+^-N always remains higher in soils treated with plantain urines compared to ryegrass, with lower nitrification rates until after day 21 maintaining the idea of nitrification delay (see Supplementary Figure 1). In spring, NH_4_^+^-N is always lower in the soils treated with plantain urines compared to ryegrass urine while the rate of nitrification only looks to be lower until day 14. Although highly speculative, lower NH_4_^+^-N in spring could be indicative of urease inhibition by metabolites within the UOF, or possibly substrate inhibition i.e. localised high concentrations of urea causing enzyme inhibition.

It should also be noted that dilution of urines to normalise total N delivered to the soil in the incubations would have diluted metabolites within the UOF that might be associated with BNI and therefore nitrification rates. This especially affects the ryegrass-derived urine, which was diluted the most (Autumn: 1.3- to 3.5-fold dilution for plantain urines versus 4.8 for ryegrass; Spring: 1.4- to 5.4-fold dilution for plantain urines versus 7.4 for ryegrass – see Supplementary Table 1). However, at least for the plantain urines, there was no obvious relationship between higher dilution and whether NO_3_^-^-N was also higher in the soil due to lower concentrations of putative inhibitory metabolites. For example, in the spring, urine derived from Cultivar D was diluted 5.4-fold yet exhibited the greatest inhibition, while urine derived from Cultivar C was undiluted and had a mid-range inhibition. Overall, the results from the spring grazing trial were quite different to those from the autumn with respect to the ranking of cultivars relative to nitrate produced, the variation in nitrate produced at each timepoint across all cultivars, and the change in nitrification rates between each timepoint. It has been suggested that this may be a function of both the winter growth and flowering characteristics of the cultivars at the time of grazing – Cultivars D and H originate from Northern European germplasm, with low winter growth and late flowering (Table 1), while Cultivars B and E, G and I, are of Mediterranean or New Zealand origin, with high winter growth and early flowering (28 days earlier in the spring grazing trial than cultivars D and H). It is possible that the higher total N concentration in the spring urines is a ‘concentration’ effect because of decreased plantain intake and urinary-N concentration due to a greater proportion of less palatable scape present at the time of grazing, particularly in the early flowering cultivars. However, there was no noticeable difference between either the bodyweight or condition of the sheep grazing the different cultivars at the end of the grazing period or the residuals in the plots after each grazing in both seasons (Judson, Agricom, *pers. comm.,* 2017).

### 4.2 Metabolomic analysis of leaf extracts as a primary source of BNI associated metabolites

Alongside relatively small changes in aucubin and catalpol, the high concentrations of verbascoside in spring compared with autumn (see Figure 4) corresponds with observations previously seen for plantain (Tamura, 2002; Navarrete et al., 2016; Box et al., 2019). This suggests verbascoside does play a role in reducing the accumulation of NO_3_^-^-N in the soil, but leaf concentrations of verbascoside would be an unreliable indicator of cultivar performance with respect to BNI. For example, the poorer incubation results for some cultivars (greater NO_3_^-^-N accumulation after 28 days) despite reasonable verbascoside concentrations, such as Cultivar C grazed in spring, compared with cultivars with much lower verbascoside and lesser NO_3_^-^-N production in the soil incubation, such as Cultivar I grazed in autumn, points to additional mechanisms being at play. More obvious in Figure 4 is the alternative intermingling of autumn and spring samples relative to nitrate concentrations in soil where spring samples have consistently high verbascoside concentrations and autumn consistently low – again suggesting BNI is linked to multiple metabolites.

In addition to the oft reported verbascoside in BNI studies, this study identified additional phenylethanoid glycosides (PhGs) that also contribute to lowered NO_3_^-^-N concentrations in the soil. These included verbascoside derivatives campneoside I (L1), ß-hydroxyverbascoside (L3) and oxoverbascoside (L4), along with closely related plantamajoside (L2) where the rhamnose moiety found in verbascoside is replaced by glucose (Fuji et al 2024, Marčetic et al 2025). Whether these additional PhGs act independently as BNI inhibitors, or act in additive or synergistic ways is unknown, but all these metabolites have known antimicrobial, antioxidant and/or anti-inflammatory activity that could contribute to inhibition of nitrifying bacteria in soil (Kang et al 1994, Ravn et al 2015, Fuji et al 2024, Stochmal et al 2024, Marčetic et al 2025). The remaining metabolites contributing to lowered NO_3_^-^-N concentrations in the soil were all iridoid glycosides (IrGs) with aucubin and it’s derivative catalpol being the most discussed in the literature (e.g. Roderguez et al 2023). This current research suggests that globularin (L7), asperuloside (L9), geniposidic acid (L11) and arborescosidic acid (L12) potentially play a synergistic role in BNI while cultivar differences look to be associated with desacetylhookerioside (L6), dumiloside (L8), mussaenosidic acid (L10), catalpol (L13) and aucubin (L14).

Nitrification inhibition has been linked with aucubin but not catalpol (Rodriguez et al., 2023 and references therein), while the other IrGs identified in this study have not been directly associated with BNI or other bioactive properties such as antimicrobial activity; rather, they have been found within suites of IrGs, that include bioactive metabolites (Jensen et al 1996, Buathong et al 2018, Chan et al 2020). If aucubin and catalpol are more associated with defining cultivar physiology as opposed to being correlated with BNI, this could provide a partial explanation for the mixed reports in the literature related to BNI effects and nitrous oxide emissions (Dietz et al., 2013, Rodriguez et al 2023, Gardiner et al., 2018 and 2020). However, it should be noted that there are significant seasonal and diurnal differences in the concentrations aucubin, catalpol and verbascoside (Box et al., 2018) with these fluctuations also validly invoked to account for mixed BNI effects. Similar seasonal and even daily fluctuations are likely for all other metabolites.

The observations presented in this study are inferences only as opposed to causal relationships directly linking plantain leaf metabolites to BNI, however, they do provide guidance as to possible plant derived metabolites that contribute to urine chemistries conducive to BNI. As outlined in section 4.5 below, of particular note are the PhG metabolites within the cohort of urinary metabolites identified that are associated with lowered NO_3_^-^-N concentrations in the soil (see Table 4).

### 4.3 Bioassay of the urine organic fraction (UOF) and the relationship to soil nitrification assays

The shape of the dose response curve (Supplementary Figure 3) was more linear than the asymptotic relationship reported in prior work by Peterson et al. (2022) and the mass of UOF added in these experiments that elicited 100% inhibition was approximately half. This suggests the proportions of metabolites within the UOF that might contribute to BNI vary considerably in concentration and are likely to be affected by many different variables. Variation in plant chemistry contributing to these differences in resulting UOF chemistry is probably related to factors that influence plant physiology such as season, temperature, plant age and nutrient supply, but there will also have been animal metabolic differences (Adler et al., 1995; Tamura, 2001; Tamura, 2002; Tamura and Nishibe, 2002; Al-Mamun et al., 2008; Navarrete et al., 2016; Box and Judson, 2018; Box et al., 2019). Despite all these variable factors, the bioassay showed a similar order of cultivars with respect to their nitrification inhibition potential when compared with the autumn soil incubations.

The ‘order of cultivars’ with respect to nitrification inhibition in the soil assay was different in spring and this study lacks a follow-up bioassay of spring UOFs. Prior to these between season cultivar dependant nitrification differences being realised, the autumn bioassay results had already been carried forward and used in the correlative discriminatory analysis for narrowing down BNI associated spring urine metabolite targets. It should be noted that this unfortunately weakens the analysis and has potentially removed some metabolic targets from consideration.

It is likely that the results derived from the bioassay provide a more robust comparison between cultivars with respect to BNI potential than the soil incubations. This is because the assay compares equal concentrations of UOF as opposed to varying dilutions of UOF in the soil incubations that resulted from normalising urines based on total N. Unexpectedly, there was no correlation between NUNC mass per litre urine and % inhibition. As mentioned above, this suggests that the proportions of metabolites within each UOF (dominated by NUNC) that contributed to BNI varied considerably in concentration and provides a partial explanation for seasonal differences and general variability in the nitrification rates observed. As such, no singular cultivar will be perfect year-round, meaning compromises must be made concerning nitrification inhibition potential and agronomic features such as winter activity and persistence in the sward.

It should also be noted that the bioassay only tested the inhibition of one ammonia-oxidising (nitrifying) bacterium, and it is known that different nitrifying microorganisms (including archaea) have differing sensitivity in the presence of the same BNI-associated plant metabolites (Kaur-Bhambra et al., 2022; Kolovou et al., 2023). Although the bioassay allows more robust comparison of different UOFs, the rates of NO_3_^-^-N production observed in soil incubation experiments will still be a more realistic gauge of what might happen in the field, where a consortium of soil microbes will be exposed to the UOF metabolites and impacts on NO_3_^-^-N production will represent whole community sensitivity.

### 4.4 Key urine metabolites with respect to plant chemistry and BNI in soil

The compounds identified as “urine metabolites” and “aromatic metabolites” in Table 4 are thought to be due to metabolic changes by the animal, or its microbiome, because of consuming plantain. Compounds U8 (diethyl-2-hydroxy-3-methylsuccinate) and U9 (2-/3-isopropylmalic acid) accumulated as a likely result of valine (for U8), leucine and isoleucine metabolism (for U9) (Strassman and Ceci, 1963). A natural origin for U8 could not be found in the literature, although it may be derived from rumen fermentation as related compounds have been reported from bovine ruminal fluid (Eom et al., 2022; Connolly et al., 2024) and reductase enzymes for precursor compounds are known from fermentative *Saccharomyces* species (Furuichi et al., 1987).

4-Hydroxy-2-oxopentanedioic acid (U12) may be microbially derived since it has been reported as an intermediate of catechol degradation yielding pyruvate via a plasmid enzyme mediated metabolic pathway in *Pseudomonas* species (Kunz et al., 1981). 4-Hydroxy-2-oxoheptanedioic acid can also be derived from pimelic acid which in turn is a product of fatty acid synthesis and is part of a biotin biosynthetic pathway in *Bacillus subtilis* (Manandhar and Cronan, 2017). A degradation enzyme for this compound is known from *Escherichia coli,* producing pyruvate and succinic semi-aldehyde (Rea et al., 2007). Although 4-Hydroxy-2-oxoheptanedioic acid has not been reported as a urinary metabolite, pimelic acid along with biotin metabolites have (Bennett et al., 1992; Peterson et al., 2004). Whether or not this compound has any influence on inhibiting nitrification or ureolysis is unknown.

Compound U10 (undefined sulfonic acid; putatively identified as (4-ethyl-2,6-dihydroxyphenyl) oxidanesulfonic acid) has been reported in rat urine but its metabolic origin was not discussed (Zhu et al., 2021), while 3-(3-Hydroxyphenyl)propanoic acid (U19) is a known mammalian urine metabolite (Booth et al., 1960) derived from bacterial metabolic degradation of aromatics, e.g. caffeic and chlorogenic acid, where these compounds are known to have antibacterial activity (Wang et al., 2022). Compound U18 (Vanillylmethanol 4-sulfate) is also likely derived from metabolism of aromatic compounds. Isovanillic acid-3-sulfate (U11), may be derived from microbial breakdown of dietary polyphenolics (Curti et al., 2025) and is also known to have antimicrobial activity (Matejczyk et al., 2024). Azelaic acid (U13) is a well-documented antibacterial agent (Sauer et al., 2023) but is also found in the rumen fluid of sheep and cows, derived from the microbial hydrogenation of non-conjugated dienoic acids (de Almeida et al., 2018). Interestingly, non-covalent synthetic derivatives combining azelaic acid and urea have been investigated to improve the N use efficiency of fertilisers by slowing urea release rates (Faulks et al., 2025).

Metabolite U17 (guaiacol-ß-D-glucuronide) correlated well with the glucuronidation of 2-methoxyphenyl (also known as guaiacol), potentially sourced from lignin and phenolic acids through bacterial fermentation in the rumen (Martin, 1982). The metabolite 2-methoxyphenyl sulfate (U16) may also be associated with this pathway where sulfation can be used as a route for excretion of hydrophilic metabolites (Correia et al., 2021; Zhuang et al., 2021). Guaiacol in wastewater from woodchip gasification has been shown to be strongly inhibitory to nitrification (Jansen, 2003). Catechol glucuronide (U15) may be a breakdown product of catecholamines (present in both animals and plants – including plantain), which are in-turn derived from tyrosine (Kuklin and Conger, 1995; Kulma and Szopa, 2007). It is interesting to note that a co-crystalline synthetic form of catechol combined with urea can act as a urease inhibitor (Casali et al., 2019), which would then affect nitrification rates. Catechol can also be released from plant roots and is known to be mildly antimicrobial (Kocaçalişkan et al., 2006; Vishakha et al., 2020).

### 4.5 BNI associated urine metabolites derived from verbascoside, aucubin and catalpol

As has been previously reported, the plantain bioactive compounds aucubin, catalpol, and verbascoside were not detected in animal urine samples (Peterson et al., 2022). However, based on mass spectral fragmentation patterns, metabolic derivatives of these compounds were observed as methylated, sulfated, glucuronated and combinations thereof in the urine (Visioli et al., 2003; Qi et al., 2012; Qi et al., 2013; Xu et al., 2023). Metabolites U1, U2, U3, U4, U5 and U6 are all derivatives of the plantain PhGs. Analysis of most of the plantain cultivars indicated that the largest discriminator of phenylethanoid origin among leaf metabolites, and the likely source of the hydroxytyrosol moieties implicated in BNI, was verbascoside, although for Cultivar G, plantamajoside also had a major influence.

Verbascoside-derived metabolites such as hydroxytyrosol and other polyphenols have been reported to scavenge nitrosating compounds under defined laboratory conditions (Kono et al., 1995; De Lucia et al., 2008). Hydroxytyrosol is also known to inhibit the expression of nitric oxide synthase (iNOS) in animal models preventing the production of nitric oxide (NO) (Maiuri et al., 2005). It is unknown whether there is a relationship between the hydroxytyrosol compounds and soil N cycling processes like co-denitrification that specifically require nitrosating compounds, NO and/or NO_2_^-^ (Spott et al., 2011; Kirkby et al., 2025). Hydroxytyrosol from olive oil byproducts has recently been valorised as a ‘sustainable’ BNI active compound against the nitrifying microorganism *Nitrosomonas europaea* (Fernández et al., 2025). While the variation in verbascoside with plant age, nutrient supply, temperature and light quality is well documented (Bowers and Stamp, 1993; Tamura, 2001; Tamura, 2002; Box et al., 2019), there is little information on how these factors influence the expression of other phenylethanoids that may also be a source of hydroxytyrosol moieties (Budzianowska et al., 2004).

Although verbascoside dominated derivative metabolite features that were associated with BNI, catalpol and derivatives potentially sourced via gut microflora metabolism (Tao et al., 2016) also may have influenced soil N cycling. The aucubigenin derivative (U7) is possibly derived from catalpol or its precursor, aucubin (Damtoft, 1994). Rodriguez et al. (2023), note that aucubigenin can inhibit cytochrome P-450 and postulate that aucubigenin’s structure might inhibit the enzyme ammonia monooxygenase (AMO) in soil resulting in inhibition of the first step in nitrification.

## Conclusions and future research

Nitrification inhibition in soil by urine metabolites from sheep grazed on nine different plantain cultivars was compared with animals fed ryegrass. Overall, these results suggest that ingestion of phenylethanoid and iridoid glycosides resulted in ovine urine secondary metabolite profiles that lowered NO_3_^-^-N in soil incubations. Metabolites derived from verbascoside such as hydroxytyrosol sulfate were the most likely contributors to lowered NO_3_^-^-N in soil, with glucuronide metabolites such as catechol and guaiacol also implicated. Based on the literature available for the urinary compounds identified, the likely mechanisms of action on the nitrifying bacterial community are both bactericidal, through direct antimicrobial action, or bacteriostatic, where key enzymes such as urease or AMO could be inhibited.

Future research requires confirmation of the presence and influence of these metabolites along with establishment of more robust BNI indicators and a more representative bioassay to unravel the complexities of the physical-chemical-biological interactions involved. Some specific areas to focus on include deciphering how plantain-derived phenylethanoids and phenolics might act as scavengers of nitrosating species, as microbial inhibitors, or other mechanisms. The importance of glucuronide metabolites also needs to be established along with the function of catalpol and its metabolites in relation to microbial interactions in the ovine digestive system and correlation with altered soil N processes under urine patches. Additional trials and analyses focused on plant physiology are also warranted to fully understand the expression of metabolites associated with BNI. Finally, *in-situ* measurements of N leaching and nitrous oxide emissions are required to specifically corroborate the effects of urine metabolites identified in this study derived from ruminants grazing plantain.

## Acknowledgements

The authors thank Loreto Hernandez, Emilie Batt, Teri Robson, Richard Gillespie, Chris Dunlop, Vanessa Hampton, Rebekah Tregurtha, Kathryn Lehto and Duncan Hedderley for their analytical and technical expertise and assistance with the experimental design. The authors also thank Paul Mudge, Kate Fransen, Ina Pinxterhuis, Penny Tricker and other collaborators for their review and comments.

This work was originally conducted under the Greener Pastures project funded by the Callaghan Innovation and PGG Wrightson Seeds. The write-up of this manuscript was supported the Sustainable Food and Fibre Futures Fund of the New Zealand Ministry for Primary Industries Plantain Potency and Practice programme (2021–2027) a DairyNZ-led, NZ-wide collaborative research and development initiative, and co-funded by NZ dairy farmers through DairyNZ Inc, PGG Wrightson Seeds Ltd. and Fonterra.

## Notes

### Competing Interest Statement

All authors reports financial support was provided by Callaghan Innovation Research Ltd, PGG Wrightson Seeds, Ministry for Primary Industries (NZ), DairyNZ, Fonterra. Glenn Judson reports a relationship with PGG Wrightson Seeds Ltd that includes: employment. The cultivars included in this work are part of a breeding programme at PGG Wrightson Seeds. This work has informed selection of cultivars taken forward for commercial development. If there are other authors, they declare that they have no known competing financial interests or personal relationships that could have appeared to influence the work reported in this paper.

### Summary of Updates

This version incorporates minor revisions to figures and figure legends.

