## Supplemental tables and figures for "Linking plantain derived metabolites in sheep urine with nitrification inhibition in soil"

**Supplementary material:**

**Supplementary Table 1:** Bulk urine N chemistry and dilution rates to normalise N addition to soil microcosms. Medium or high winter activity cultivars are highlighted in grey.

| Cultivar | Autumn urine bulk | | | | | | Spring urine bulk | | | | | |
|---|---|---|---|---|---|---|---|---|---|---|---|---|
| | Total N (mg N $L^{-1}$) | $NH_4^+$-N (mg N $L^{-1}$) | Urea-N (mg N $L^{-1}$) | NUNC† (mg N $L^{-1}$) | Soil-$NO_3^-$ (mg $kg^{-1}$) | Dilution (fold) | Total N (mg N $L^{-1}$) | $NH_4^+$-N (mg N $L^{-1}$) | Urea-N (mg N $L^{-1}$) | NUNC† (mg N $L^{-1}$) | Soil-$NO_3^-$ (mg $kg^{-1}$) | Dilution (fold) |
| **A** | 1625 | 23 | 1530 | 95 | 275 | 1.7 | 3410 | 37 | 2717 | 693 | 232 | 2.7 |
| **B** | 1330 | 22 | 756 | 574 | 261 | 1.4 | 5170 | 58 | 4090 | 1080 | 250 | 4.1 |
| **C** | 3070 | 44 | 2213 | 857 | 238 | 3.2 | 1260 | 18 | 868 | 392 | 260 | 1.0 |
| **D** | 2060 | 81 | 1541 | 519 | 237 | 2.1 | 1990 | 117 | 1429 | 561 | 189 | 1.6 |
| **E** | 1660 | 83 | 1317 | 343 | 248 | 1.7 | 6760 | 55 | 6078 | 682 | 266 | 5.4 |
| **F** | 1280 | 31 | 644 | 636 | 253 | 1.3 | 4610 | 39 | 3782 | 828 | 248 | 3.7 |
| **G** | 974 | 24 | 504 | 470 | 247 | 1.0 | 5010 | 46 | 4286 | 724 | 231 | 4.0 |
| **H** | 3390 | 71 | 2437 | 953 | 231 | 3.5 | 1730 | 71 | 1261 | 469 | 215 | 1.4 |
| **I** | 1810 | 75 | 1373 | 437 | 201 | 1.9 | 5370 | 68 | 4482 | 888 | 246 | 4.3 |
| **Ryegrass*** | 4660 | 484 | 4174 | 486 | 270 | 4.8 | 9370 | 76 | 8627 | 743 | 288 | 7.4 |

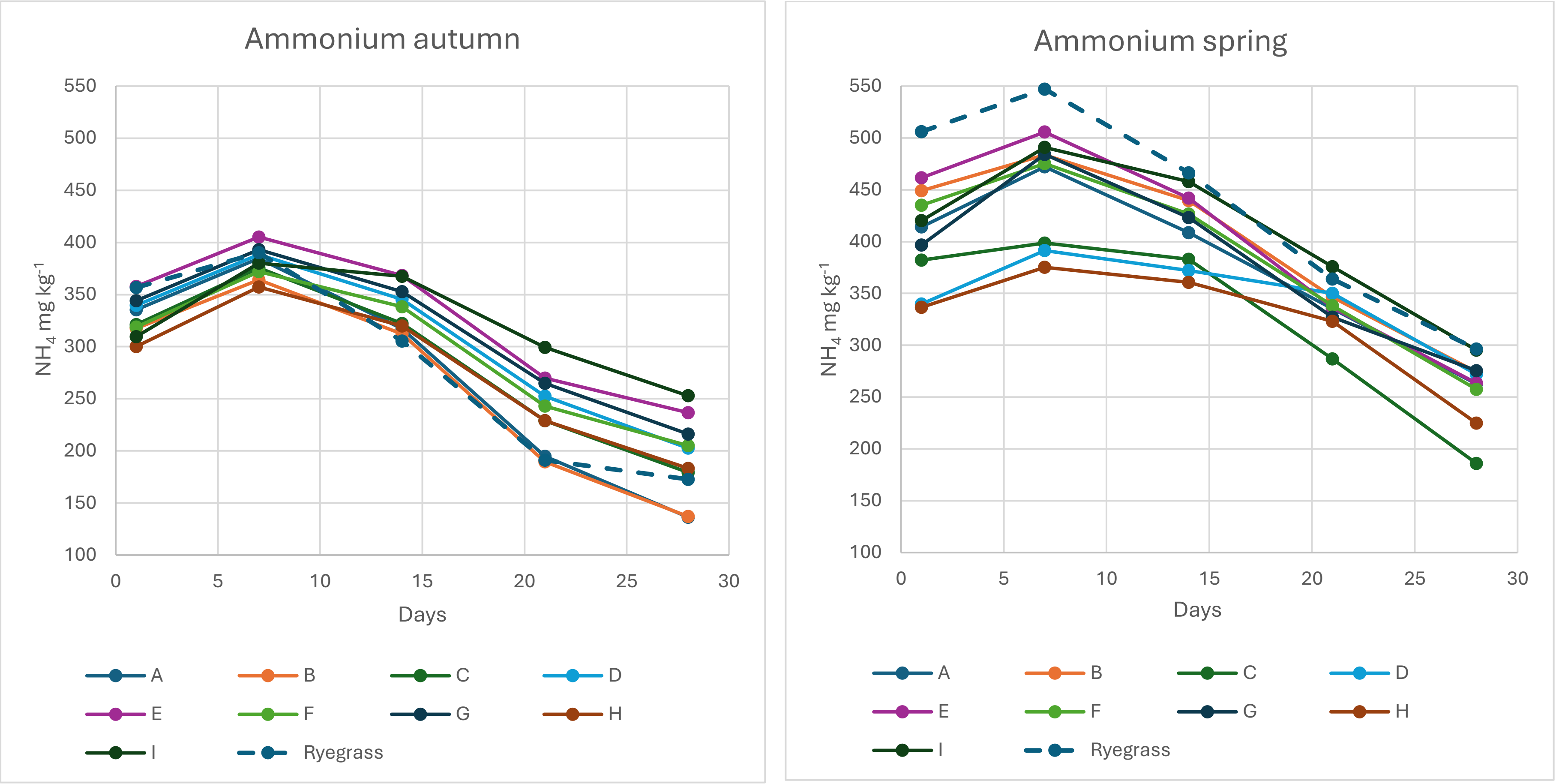


**Supplementary Figure 1:** Ammonium concentrations (means) recorded with time in the urine nitrification soil assay microcosms in autumn and spring. Ammonium in the no-urine control was not measured.

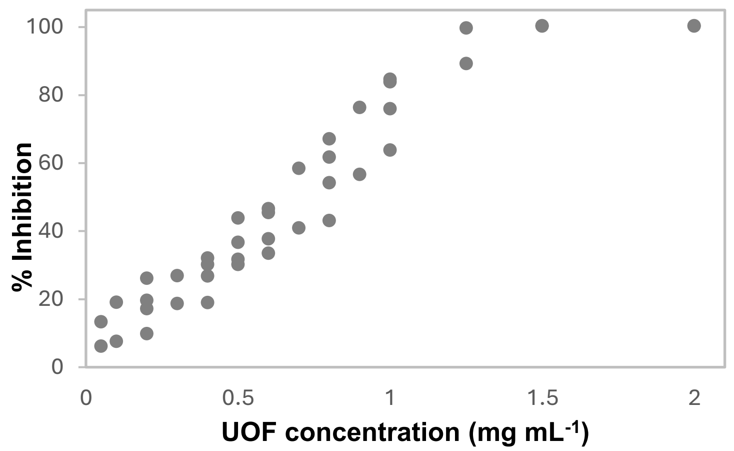


**Supplementary Figure 2:** Biological nitrification inhibition (BNI) dose response curve for the urinary organic fraction (UOF) derived from sheep fed 100% Cultivar G. Root exudates from this cultivar exhibited moderate to high BNI in prior hydroponic experiments (data not shown), moderate reduction in $NO_3^-$-N production in soil incubations in both autumn and spring (this work), had good winter activity from an agronomic perspective (as outlined in Table 1), and is a key cultivar within New Zealand plantain breeding programmes.

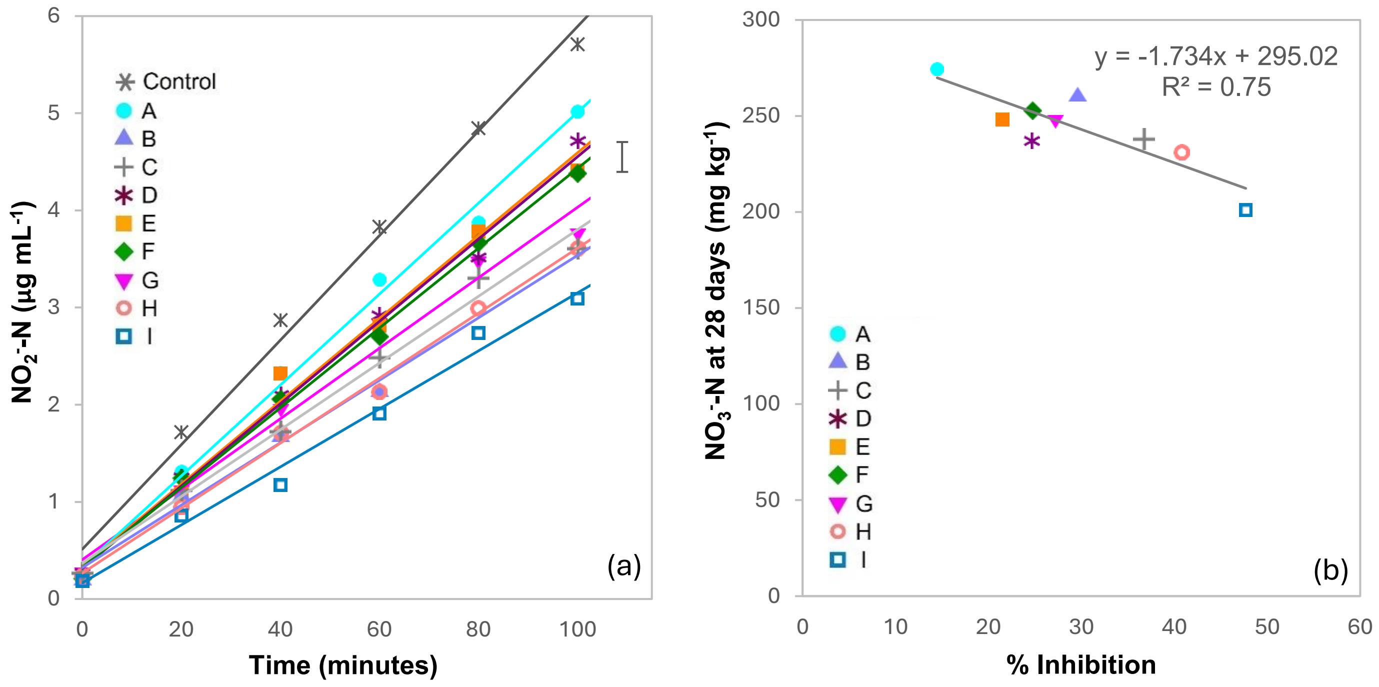


**Supplementary Figure 3:** Nitrite produced when the urinary organic fraction (UOF) (0.5 mg mL$^{-1}$) containing the non-urea nitrogenous compounds (NUNC) was assayed against a culture of the ammonia oxidising bacterium *Nitrosospira multiformis* (a). Urine was derived from sheep fed 100% plantain (Cultivars A–I) from the autumn trial. % inhibition of nitrification relative to the uninhibited (non-UOF) control plotted against nitrate produced in soil microcosms after 28 days of incubation with the same urines tested in the bioassay (b). Inhibition caused by the ryegrass urine was excluded from the analysis due to excessive colouration in the sample interfering across wavelengths measured within the colorimetric assay. The error bar in (a) represents post-hoc $LSD_{5\%}$ comparing the regressions. Figure 4a is a permitted reproduction of data published in the IGC Proceedings 2023, Peterson et al. (2023).

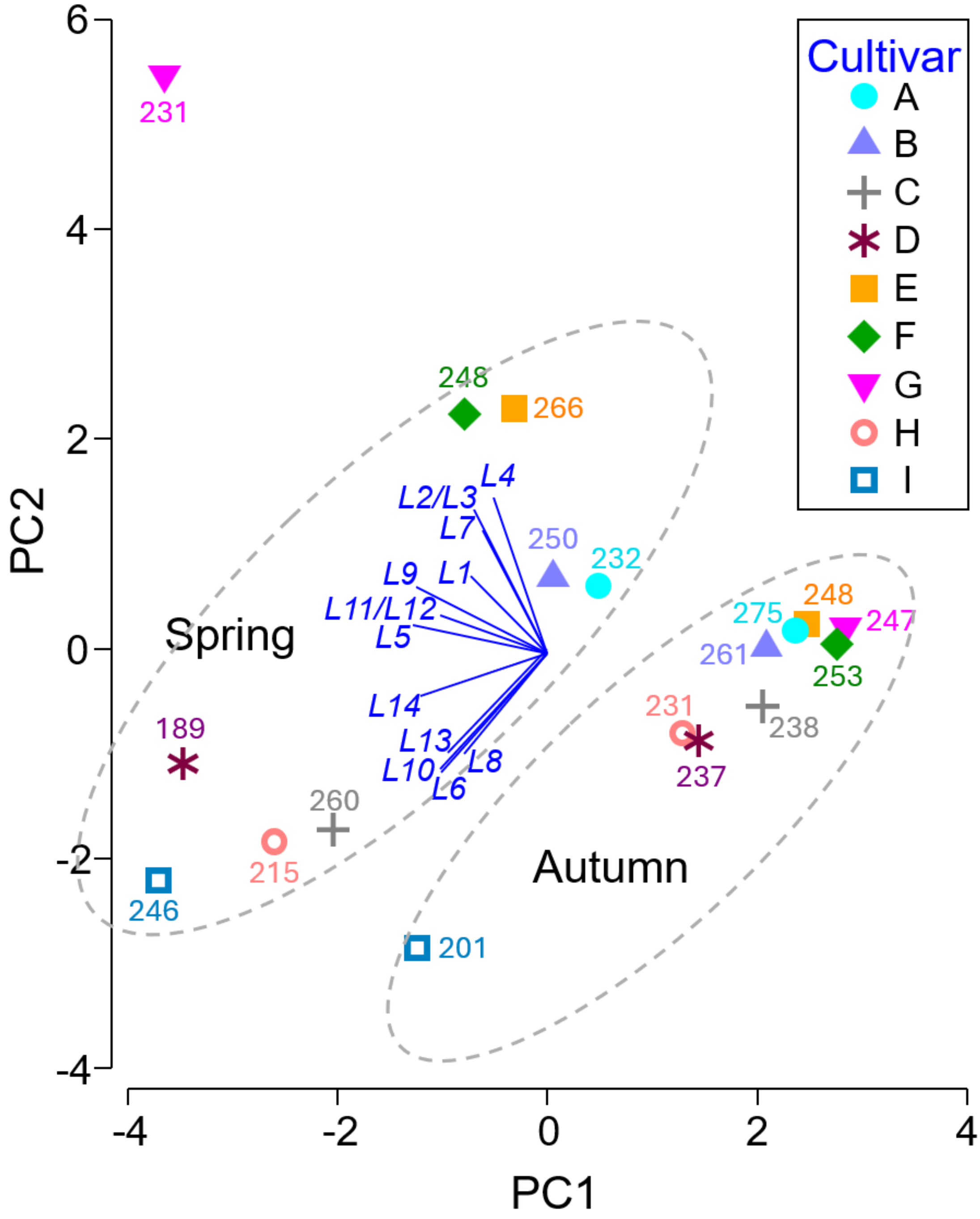


**Supplementary Figure 4** Principal components analysis (PCA) of the relative abundance of metabolites L1 to L14 presented in Table 3 identified within the leaf of plantain Cultivars A–I in the autumn and spring grazing trials. Dotted ovals are not clusters, rather they are an annotation to denote which data points belong to which season. The PC1 axis captures 45.3% of variation with a further 31.1% captured by the PC2 axis. The PC3 axis resolved a further 9.9% of variation suggesting the 2-dimensional representation here is adequate. Numbers associated with each data point are the concentrations of nitrate (mg $kg^{-1}$) measured in the soil microcosms after 28 days of incubation with urine from groups of animals fed one of the nine different plantain cultivars. The blue vector fan indicates increasing normalised area (greater concentration) of metabolites across both seasons from the centre out, with length and the direction being an indicator of 'influence' that each metabolite has with respect to where data points are positioned in the plot relative to each other.
